# Revisiting genomes of non-model species with long reads yields new insights into their biology and evolution

**DOI:** 10.1101/2023.10.06.561169

**Authors:** Nadège Guiglielmoni, Laura I. Villegas, Joseph Kirangwa, Philipp H. Schiffer

## Abstract

High-quality genomes obtained using long-read data allow not only for a better understanding of heterozygosity levels, repeat content, and more accurate gene annotation, and prediction when compared to those obtained with short-read technologies, but also allow to understand haplotype divergence. Advances in long-read sequencing technologies in the last years have made it possible to produce such high-quality assemblies for non-model organisms. This allows us to revisit genomes, which have been problematic to scaffold to chromosome-scale with previous generations of data, and assembly software. Nematoda, one of the most diverse, and speciose animal phyla within metazoans, remains poorly studied, and many previously assembled genomes are fragmented. Using long reads obtained with Nanopore R10.4.1 and PacBio HiFi, we generated highly contiguous assemblies of a diploid nematode of the Mermithidae family, for which no closely related genomes are available to date, as well as a collapsed assembly and a phased assembly for a triploid nematode from the Panagrolaimidae family. Both genomes had been analysed before, but the fragmented assemblies had scaffold sizes comparable to the length of long reads prior to assembly. Our new assemblies illustrate how long-read technologies allow for a much better representation of species genomes. We are now able to conduct more accurate downstream assays based on more complete gene and transposable element predictions.

## 1 Introduction

Over the past decade, the field of genome assembly has experienced major improvements fueled by the development of high throughput sequencing techniques and major increases in the length and accuracy of reads. Short-read sequencing prompted the release of many draft assemblies for a large variety of species. The limited length of these reads could only yield highly fragmented assemblies, which were sufficient for initial analyses of gene content, but could not account for the structure of genomes and often fell short on resolving repetitive regions [1]. Recent advances in genome assembly have been driven by the availability of long reads offered by Pacific Biosciences (PacBio) and Oxford Nanopore. While these reads initially had a high error rate, newer technologies have drastically increased their accuracy to over 99%, with the release of PacBio HiFi reads (based on circular consensus sequencing) [2] and Nanopore Q20+ [3] reads obtained using R10.4.1 flow cells. These developments have brought draft assemblies to Megabase-level N50s [4], illustrating their high contiguity, and have opened new possibilities for genome analyses. Assemblies obtained with long-read data not only have a higher gene completeness, but they can also provide a more comprehensive overview of repetitive regions and potentially allow for a better understanding of their structure, activity and dynamics [5]. In addition, high-accuracy long reads can be used to discriminate alleles and generate phased assemblies, including all haplotypes [6, 7], while low-accuracy long reads were only sufficient for collapsed assemblies (in which homologous chromosomes are represented by a single sequence) as errors could not be distinguished from alternative haplotypes.

Some hundreds of genome assemblies have been released this far for the phylum Nematoda, yet only a few are high-quality assemblies and they offer a poor representation of the diversity of the clade for which over 30,000 species have been described [8]. In particular, efforts have focused on *Caenorhabditis* and other parasitic species, leaving scarce resources for understudied clades [9, 10]. In this paper, we focus on the genomes of two species at two extremities of the nematode phylogeny: the basal *Romanomermis culicivorax* (clade I) and the derived *Panagrolaimus* sp. PS1159 (clade IV).

The Enoplean nematode *Romanomermis culicivorax* is a member of the mermithidae family which includes over 100 described species [11]. It is an obligate parasite of various species of mosquito larvae [12]. Along other mermithid nematodes it is presently employed for the biological control of malaria [13, 14]. Enoplean research often revolves around *Trichinella spiralis*, given its significance as a mammalian parasite [15]. Among mermithidae, only two genomes are currently publicly available [16, 17]. In contrast, the long-read genome assembly of the parthenogenetic *Mermis nigrescens* is more contiguous and contains approximately twice the repeat content and heterozygosity of the sexual *R. culicivorax*. The need for additional high-quality genomes is evident, not only to address resource gaps in the Enoplean class, but also to enable investigations into sexual evolution, genome structural variations, and host-parasite interactions within the mermithidae family.

*Panagrolaimus sp.* PS1159 is a free-living nematode belonging to the Panagrolaimidae family. Members of this family have various reproductive modes including hermaphroditism, outcrossing between males and females and asexual reproduction through parthenogenesis [18]; *Panagrolaimus* sp. PS1159 is parthenogenetic. This strain has been isolated in North Carolina, USA by Paul Sterneberg, and is thought to be a triploid allopolyploid (3n=12) [19]. Previous studies have found it shares a common origin of parthenogenesis with most *Panagrolaimus* asexual strains, from a hybridization event estimated to have occurred 1.3–8.5 Million years ago [19, 20]. To date, over 140 strains of the genus have been documented (NCBI Taxonomy Browser), however only nine, largely fragmented, draft genome assemblies are available on GenBank (accessed on 06.10.2023). This widely distributed group includes strains isolated from extreme environments such as Antarctica, the volcanic island of Surtsey and the Russian permafrost. Representatives of the genus from these locations have been found to be freezing-tolerant undergoing cryptobiosis [20, 21], and *P.* sp. PS1159 has also shown anhydrobiotic potential as a fast desiccation strategist [22].

Short-read genome assemblies are available for both species, yet their high fragmentation impedes downstream analyses. Their scaffold N50s are limited to 17.6 kb for *R. culicivorax* and 9.9 kb for *Panagrolaimus* sp. PS1159. By contrast, these values would be the expected length for unassembled long reads nowadays. Although these draft assemblies provided a first insight into the genomics of these species, more contiguous assemblies can now be obtained using long reads. To reassemble these species, we chose to generate both PacBio HiFi and Nanopore sequencing data and to leverage distinct advantages of these technologies. For PacBio HiFi, we used an ultra-low input protocol with DNA extracted from only a few individuals and whole genome amplification. For Nanopore sequencing, we extracted DNA from large pools of individuals and selected the largest fragments. Using these heterogeneous long-read datasets, we produced new highly contiguous assemblies with increased completeness.

## 2 Material & Methods

### 2.1 Pacific Biosciences HiFi sequencing

Up to 10 individuals were collected and washed in water, then flash-frozen using liquid nitrogen in a salt-based extraction buffer (Tris-HCl 100 mM, ethylenediaminetetraacetic acid 50 mM, NaCl 0.5 M and sodium dodecylsulfate 1%). Samples were incubated overnight at 50°C after addition of 5 *μ*L of proteinase K. DNA was precipitated using NaCl 5 M, yeast tRNA and isopropanol, and incubated at room temperature for 30 minutes, then pelleted at 18,000 g for 20 minutes (4°C). The DNA was washed twice with 80% ethanol and spinned at 18,000 g for 10 min (4°C). The DNA pellet was eluted in elution buffer (D3004-4-10 Zymo Research) and incubated at 50°C for 10 minutes. RNA was removed by incubating with RNAse (Qiagen, 19101) for 1 hour at (37°C). DNA concentrations were quantified using a Qubit 4 fluoremeter with 1X dsDNA kit. HiFi libraries were prepared with the Express 2.0 Template kit (Pacific Biosciences, Menlo Park, CA, USA) and sequenced on a Sequel II/Sequel IIe instrument with 30 hours movie time. HiFi reads were generated using SMRT Link (v10, Pacific Biosciences, Menlo Park, CA, USA) with default parameters. Sequencing results are presented in Table S1.

### 2.2 Nanopore sequencing

*Romanomermis culicivorax* worms were picked from moss material supplied by Prof. Dr Edward Platzer at University of California Riverside. *Panagrolaimus* sp. PS1159 worms (isolate from North Carolina, USA) were harvested from agar plates with water and pelleted at 5,000 g for 5 min. The *P.* sp. PS1159 pellet was resuspended in a 1 M sucrose solution used for bacterial decontamination (sucrose flotation). The sample was centrifuged at 1,000 g for 3 minutes, the upper 1 mL of the supernatant containing the live clean worms was transferred to a new tube and diluted with nuclease-free water. The worms where pelleted again at 5,000 g for 5 min for further processing. Samples were incubated in cetyltrimethylammonium bromide (CTAB) buffer (polyvinylpyrrolidone 2%, Tris-HCl 100 mM, ethylenediaminetetraacetic acid 25 mM, NaCl 2 M, CTAB 2%) supplemented with 25 *μ*L of proteinase K for 1 hour (*P.* sp PS1159) or 2 hours (*R. culicivorax*). Extracts were purified with phenol-chloroform-isoamyl alcohol 25:24:1, chloroform-isoamyl alcohol 24:1, centrifugation at 16,000 g for 10 minutes (room temperature) and AMPure XP beads (Agencourt). DNA was then incubated with RNAse cocktail enzyme mix (Thermo Fischer, AM2286) for 1 hour at 37°C. DNA was fragmented in a 2 mL low-bind round bottom Eppendorf tube using a sterile 3 mm borosilicate bead (Z143928-1EA Merck) by vortexing for 1 minute at maximum speed as described in [23]. Short fragments were removed using the Short Reads Eliminator (SRE) (Circulomics, Pacific Biosciences). The DNA samples were incubated with SRE buffer for 1 hour (50°C), then the long fragments of DNA were pelleted at 10,000 g for 30 minutes (room temperature) and re-suspended in elution buffer. DNA concentrations were quantified using a Qubit 4 fluoremeter with 1X dsDNA kit.

Nanopore libraries were prepared using the Ligation Sequencing Kit LSK114 (Oxford Nanopore Technologies). The *Romanomermis culicivorax* library was loaded a first time on one R10.4 MinION flowcell. The library was recovered from the flowcell and reloaded after nuclease flush. The *Panagrolaimus* sp. PS1159 library was loaded 4 times (with nuclease flushes and fresh library loads) on one R10.4 MinION flowcell. Fast5 files were converted to Pod5 using pod5 v0.2.2. Basecalling was performed using Dorado v0.3.1 [24] in duplex mode with model dna r10.4.1 e8.2 400bps supv4.2.0 and the reads were converted to fastq using SAMtools v1.6 [25] with the module `samtools fastq`. This resulted in 5.7 Gb of Nanopore reads for *R. culicivorax* (N50: 15.9 kb) and 10.7 Gb for *P.* sp. PS1159 (N50: 33.4 kb) (Table S2). Adapters were trimmed using chopper v0.5.0 [26] with minimum quality `-q` set to default (for *R. culicivorax* and *P.* sp. PS1159) or 20 (for *P.* sp. PS1159).

### 2.3 RNA sequencing

RNA was extracted from *R. culicivorax* adults using a modified version of the protocol established by Chomczynski and Sacchi [27]. Tissue pellets of approximately 10 mg were transfered into 1 mL Trimix and lysed using a homogeniser (Ultra-Turrax, IKA Werke GmbH) for 10 minutes on ice. After addition of 200 *μ*L chloroform and incubation at room temperature for 5 min, the sample was centrifuged for 10 min at 15,000 g. The aqueous phase was collected and supplemented with 0.025 volumes of 1 M acidic acid and 0.5 volumes of pre-cooled 100% EtOH (*−* 20°C). RNA was precipitated overnight at *−* 20°C and then centrifuged at 15,000 g for 20 min. After removing the supernatant, the RNA pellet was dried for 10 min and resuspended in 125 *μ*L of GU-mix and added 3.125 *μ*L 1M acidic acid, vortexed the sample and added 70 *μ*L 100% EtOH. RNA was precipitated overnight at *−* 20°C and then centrifuged 15,000 g for 20 min and washed twice with 500 *μ*L EtOH (70%).The RNA pellet was resuspended in 20 *μ*L DEPC-H2O and incubated at 65°C for 5 min. The quality of the total RNA was assessed using degenerative agarose-gel electrophoresis and a Nanodrop 1000 photometer (Agilent Inc.). RNA libraries were prepared using a TrueSeq RNA Sample Prep kit v2 (Illumina Inc.) and sequenced on Illumina HiSeq and MiSeq platforms (Illumina Inc.) at the Cologne Center for Genomics (CCG, Cologne, Germany). For *Panagrolaimus* sp. PS1159, publicly available Illumina RNA sequencing reads were used (SRR5253560) [19].

### 2.4 Long-read preliminary analyses

Quality and length of PacBio HiFi and Nanopore reads were plotted using Nanoplot v1.41.3 [26]. Ploidy was estimated using Smudgeplot v0.2.2 [28] with the PacBio HiFi reads.

### 2.5 *Romanomermis culicivorax* long-read assembly

PacBio HiFi reads were assembled using Flye v2.9 [29] with parameter `--pacbio-hifi`, hifiasm v0.19 [6] with parameter-l 3, NextDenovo v2.5 [30] with parameters `genome size=300m read type=hifi`, and wtdbg2 v2.5 [31] with parameter `-x` ccs. For Nanopore reads, Canu v2.2 [32] was run with parameters `-nanopore genomeSize=300m`, Flye v2.9 [29] with parameter `--nano-hq`, NextDenovo v2.5 [30] with parameters `genome size=300m read type=raw`, and wtdbg2 v2.5 [31] with parameter `-x` ont. To combine PacBio HiFi and Nanopore reads, Nanopore reads longer than 15 kb were selected using seqtk v1.3 [33] with the module `seqtk seq` and the parameter `-L 15000`. hifiasm v0.19 was run using the PacBio HiFi reads and Nanopore reads *>* 15 kb with parameter `-l` 3. Assembly using Verkko v1.4 with default parameters failed.

### 2.6 *Panagrolaimus* sp. PS1159 long-read assembly

PacBio HiFi reads were assembled using Flye v2.9 [29] with parameter `--pacbio-hifi` and with the option `--keep-haplotypes`, hifiasm v0.19 [6] was run with parameters `--n-hap` 3 and `-l` set to 0 and 3, NextDenovo v2.5 [30] with parameters `genome size=300m read type=hifi`, and wtdbg2 v2.5 [31] with parameter `-x ccs`. Nanopore reads with a quality higher than Q20 were selected using chopper. Different parameters were tested to adapt to the high accuracy and assemblies with highest contiguity and completeness were selected. Canu v2.2 [32] was run with parameters `-nanopore -corrected genomeSize=300m`. Flye v2.9 [29] was run with parameters `--nano-corr` and with the option `--keep-haplotypes`. NextDenovo v2.5 [30] was run with parameters genome size=300m read type=hifi. wtdbg2 v2.5 [31] was run with parameter `-x` ont. To combine PacBio HiFi and Nanopore reads, Nanopore reads longer than 30 kb were selected using seqtk v1.3 [33] with the module seqtk seqand the parameter-L 30000. hifiasm v0.19 [6] was run using both datasets with parameters *--n-hap 3* and `-l` set to 0 and 3. Verkko v1.4 [7] was run with default parameters.

### 2.7 Assembly evaluation and post-processing

Assembly statistics were calculated using assembly-stats v1.0.1 [34]. Ortholog completeness was computed using the Benchmarking Universal Single-Copy Orthologs (BUSCO) [35] tool v5.4.7 with parameter `-m genome` against the Metazoa odb10 and Nematoda odb10 lineages. PacBio HiFi reads were mapped against HiFi assemblies using minimap2 v2.24 [36] with parameters `-ax map-hifi` and Nanopore reads were mapped against the Nanopore and hybrid assemblies with parameters `-ax map-ont`. Mapped reads were sorted using SAMtools v1.6 with the module `samtools` sort. Contigs were aligned against the nt database using BLAST v2.13.0 [37]. The outputs were provided as input to BlobToolKit v4.1.5 [38], and contaminants were subsequently removed. Reads were mapped again using minimap2 v2.24 and the output was provided to purge dups v1.2.5 [39] to remove uncollapsed haplotypes. PacBio HiFi reads were used to purge HiFi-based assemblies, Nanopore reads for Nanopore-based assemblies, Nanopore reads for hybrid assemblies of *Panagrolaimus* sp. PS1159, and PacBio HiFi reads for hybrid assemblies of *Romanomermis culicivorax* (due to the low coverage of Nanopore reads).

### 2.8 Final scaffolding

*Romanomermis culicivorax* was assembled following two pipelines: (1) the decontaminated NextDenovo PacBio HiFi contigs were purged once using purge dups; (2) the decontamined hifiasm PacBio HiFi + Nanopore contigs were purged twice; the assembly (1) was then scaffolded using RagTag v2.1.0 [40] and the assembly (2) as reference. *Panagrolaimus* sp. PS1159 was also assembled using two pipelines: (1) the decontaminated hifiasm `-l` 3 PacBio HiFi + Nanopore contigs were purged twice using purge dups; (2) the decontaminated Flye `--keep-haplotypes` Nanopore contigs were purged twice; the assembly (1) was then scaffolded using RagTag v2.1.0 and the assembly (2) as reference. The decontaminated hifiasm `-l 0` PacBio HiFi + Nanopore contigs were retained as a phased assembly.

### 2.9 Repeat and gene annotation

Repeats were annotated using the Extensive *De novo* TE Annotator (EDTA) pipeline v2.0.1 [41] with parameters `--sensitive 1 --anno 1`. This pipeline filters and combines predictions from LTRharvest [42], LTR FINDER [43] LTR retriever [44], HelitronScanner [45], Generic Repeat Finder [46], TIR-learner [47] and produces afinal transposable element library using RepeatModeler [48]. The output hardmasked assembly was converted into a softmasked assembly. RNA-seq reads were trimmed using Trim Galore v0.6.10 and mapped to the assemblies using hisat2 v2.2.1 [49]. After sorting using SAMTools v1.6 [25], the mapped reads were provided as input to BRAKER v3.0.3 [50] with parameters `--gff3 --UTR` off.

### 2.10 Downstream analyses

BUSCO v5.4.7 [35] was run on the annotated protein-coding genes using the option `-m proteins` against the Metazoa odb10 and Nematoda odb10 lineages. *k* -mer completeness of the assemblies was assessed based on the PacBio HiFi dataset using Merqury v1.3 [51].

## 3 Results

### 3.1 Initial long-read analyses

PacBio HiFi sequencing resulted in 37.5 Gb of reads (N50: 12.6 kb) for *Romanomermis culicivorax* and 29.2 Gb (N50: 15.8 kb) for *Panagrolaimus* sp. PS1159 (Table S1). Nanopore sequencing yielded 5.7 Gb (N50: 15.9 kb) for *R. culicivorax* and 10.7 Gb (N50: 33.4 kb) for *P.* sp. PS1159 (Table S2). While PacBio HiFi reads have a higher quality, Nanopore reads reach longer lengths, including some reads of 100+ kb (Figure 1a). Ploidy analyses using Smudgeplot predicts *R. culicivorax* as a diploid genome, while *P.* sp. PS1159 is expected to be triploid. Nanopore reads with Q20+ quality were selected for initial assembly of *P.* sp. PS1159, but no quality threshold was applied for *R. culicivorax* Nanopore reads due to their limited amount. All PacBio HiFi reads were used for initial assemblies.

**Figure 1:**
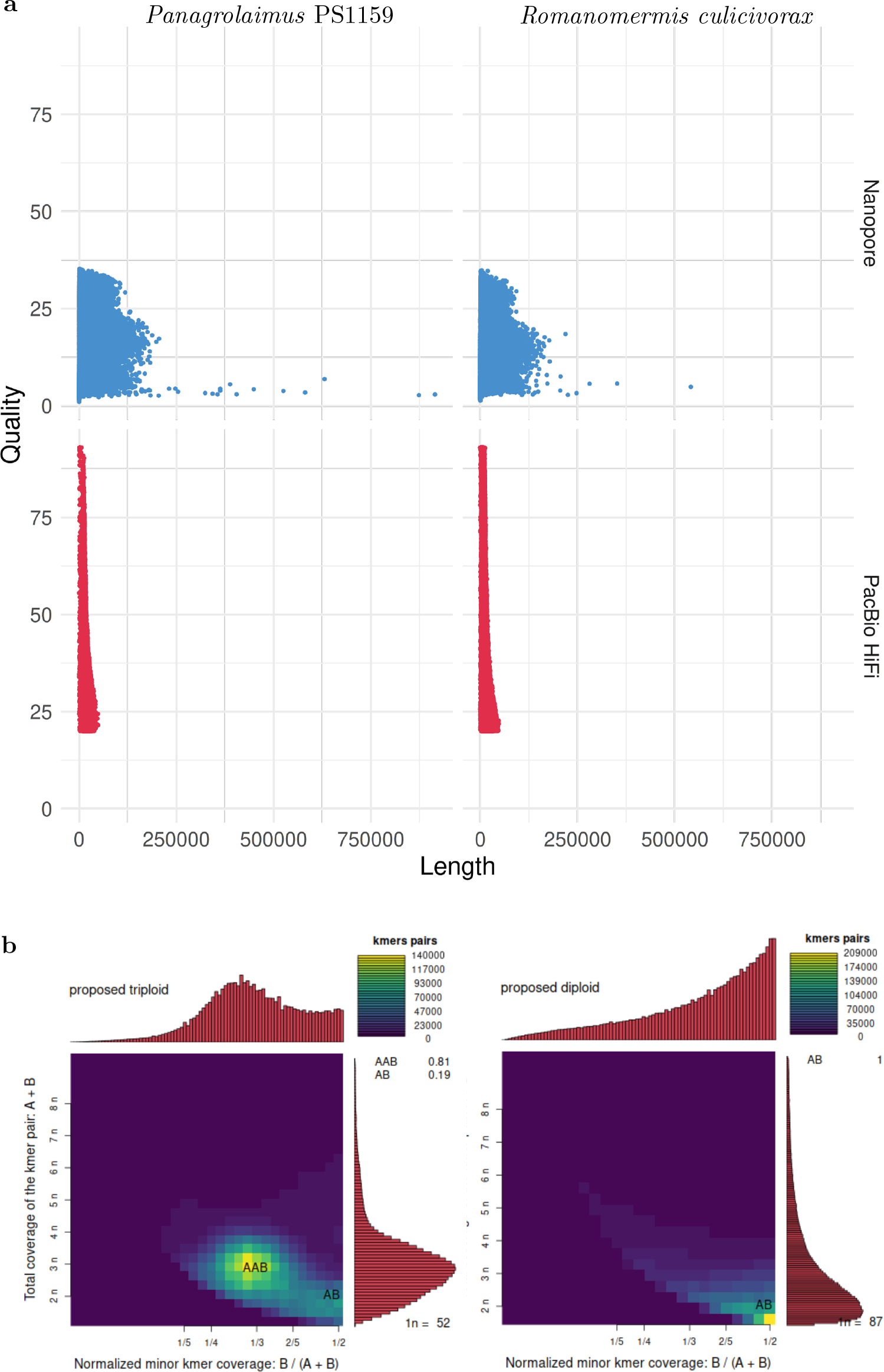
Initial analyses of the long reads. a) Quality and length of Nanopore and PacBio HiFi reads. b) *k* -mer analysis of the ploidy of the genomes using PacBio HiFi reads.

**Figure 2:**
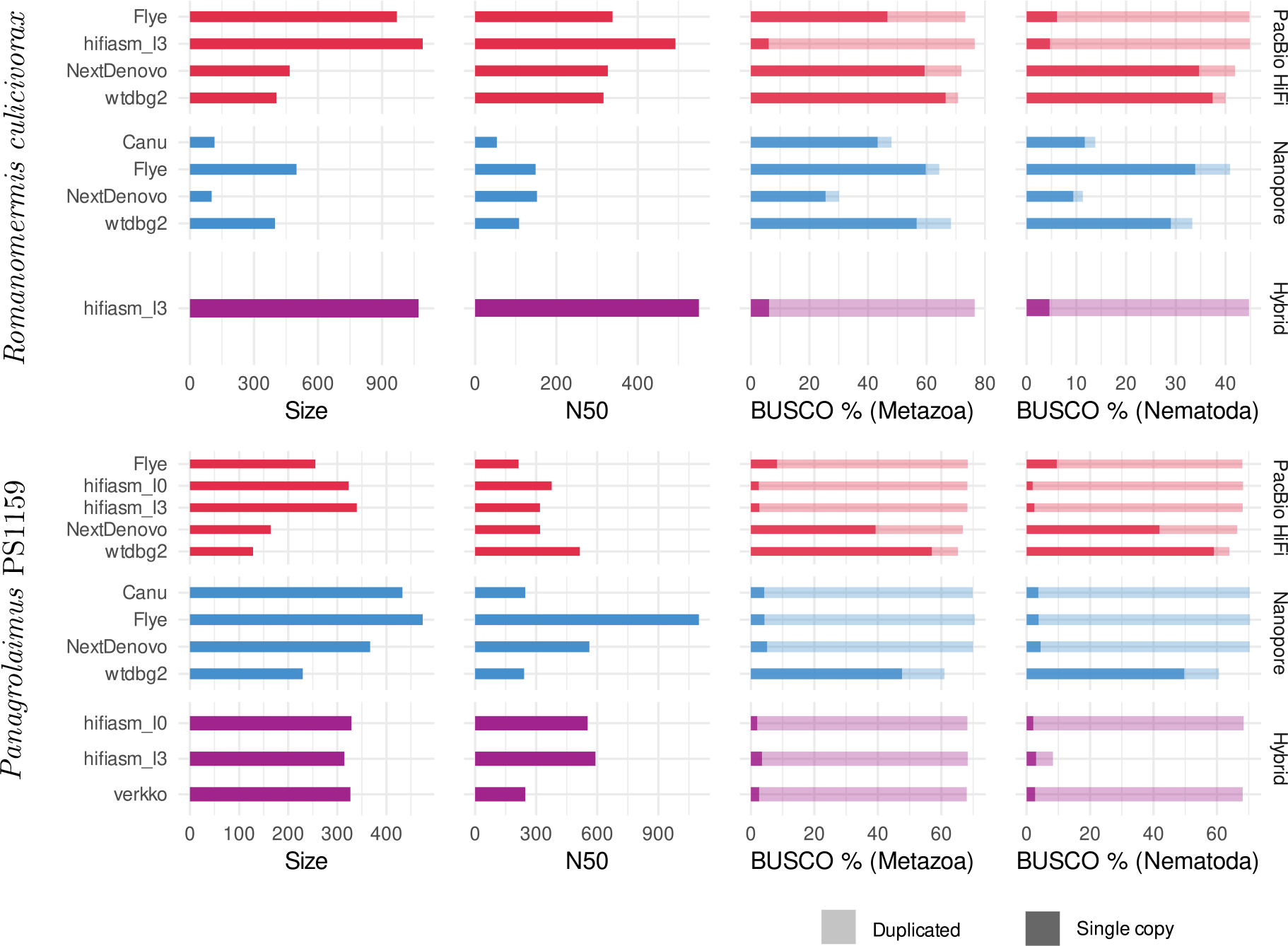
Draft assembly statistics of PacBio HiFi reads (red), Nanopore reads (blue) and the two combined (purple) with assembly size (in Mb), N50 (in kb) and BUSCO completeness against the Metazoa and Nematoda lineages.

### 3.2 High-quality long-read assemblies

Depending on the program used, assemblies of PacBio HiFi, and Nanopore reads yielded contigs with variable contiguity, and cumulative size. For *Romanomermis culicivorax*, some PacBio HiFi assemblies had a size moderately above the Illumina assembly size of 322.8 Mb (467.2 for NextDenovo, 404.8 Mb for wtdbg2), but hifiasm and Flye produced oversized assemblies (969.3 Mb and 1.1 Gb respectively). These large genome sizes could not be explained by bacterial contamination in the data coming from their environment, as there was almost none in HiFi assemblies (Figure S1). Nanopore assemblies were smaller: Flye and wtdbg2 assembly sizes were above the Illumina assembly size (499.3 Mb and 398.2 Mb), and Canu and NextDenovo assemblies were much shorter (114.9 Mb and 101.7 Mb). This is likely due to the low coverage of the Nanopore dataset, which was aggravated by a high amount of contamination from Proteobacteria and Bacteroidetes (Figure S2), and led to a suboptimal sequencing coverage for these assemblers. Therefore, it is expected for Flye and wtdbg2 to yield the most qualitative assemblies as they have been shown to be more robust with low-coverage datasets [52]. The hybrid assembly obtained using hifiasm is oversized (1.1 Gb), similar to the PacBio-HiFi-only hifiasm assembly. N50s ranged from 108 kb (wtdbg2, Nanopore) to 550 kb (hifiasm, hybrid); although these values do not reach the Megabase level, they are still one order of magnitude larger than for the Illumina assembly (17.6 kb).

For *Panagrolaimus* sp. PS1159, assemblies ranged from 128.4 Mb (wtdbg2, PacBio HiFi) to 473.9 Mb (Flye, Nanopore). Shorter assemblies correlated with a low number of duplicated BUSCO orthologs, suggesting that they would be collapsed assemblies, in which homologous chromosomes are represented by one sequence. Larger assemblies have a high number of duplicated BUSCO orthologs, indicating that haplotypes are separated. These values would match the expectation of a phased assembly with a size three times larger than a collapsed assembly, for a triploid genome. These draft assemblies were overall more contiguous than for *R. culicivorax*, with a minimum of 240 kb (wtdbg2, Nanopore) and a maximum of 1.1 Mb (Flye, Nanopore). In addition, Nanopore assemblies had fewer bacterial contaminants than PacBio HiFi assemblies (Figures S3-S4), likely owed to the supplementary sucrose decontamination step during library preparation. These read sets overall suffered much less from bacterial contamination than the Illumina data used in [19].

After decontamination, haplotig purging and scaffolding, high-quality assemblies were obtained for both species. Although long reads were not sufficient to reach chromosome level, the final assemblies had an N50 over 1 Mb (1.1 Mb for *R. culicivorax* and 3.1 Mb for *Panagrolaimus* sp. PS1159) and their contiguity is drastically improved compared to Illumina assemblies (Table 1). Furthermore, their BUSCO scores against the Metazoa and Nematoda lineages were also improved. Interestingly, the nematode BUSCO score of *R. culicivorax* remained low (35.2%), despite a higher metazoan BUSCO score. This suggests that the genome could be lacking many orthologs that would be expected in nematodes. The assembly of *R. culicivorax* has a QV score of 54.97; the *k* -mer spectrum shows a mostly collapsed assembly with yet some remaining artefactual duplications (Figure S5). The assembly of *P.* sp. PS1159 has a QV score of 47.73 and the *k* -mer spectrum also supports a mostly collapsed assembly with limited artefactual duplications (Figure S6).

**Table 1:**
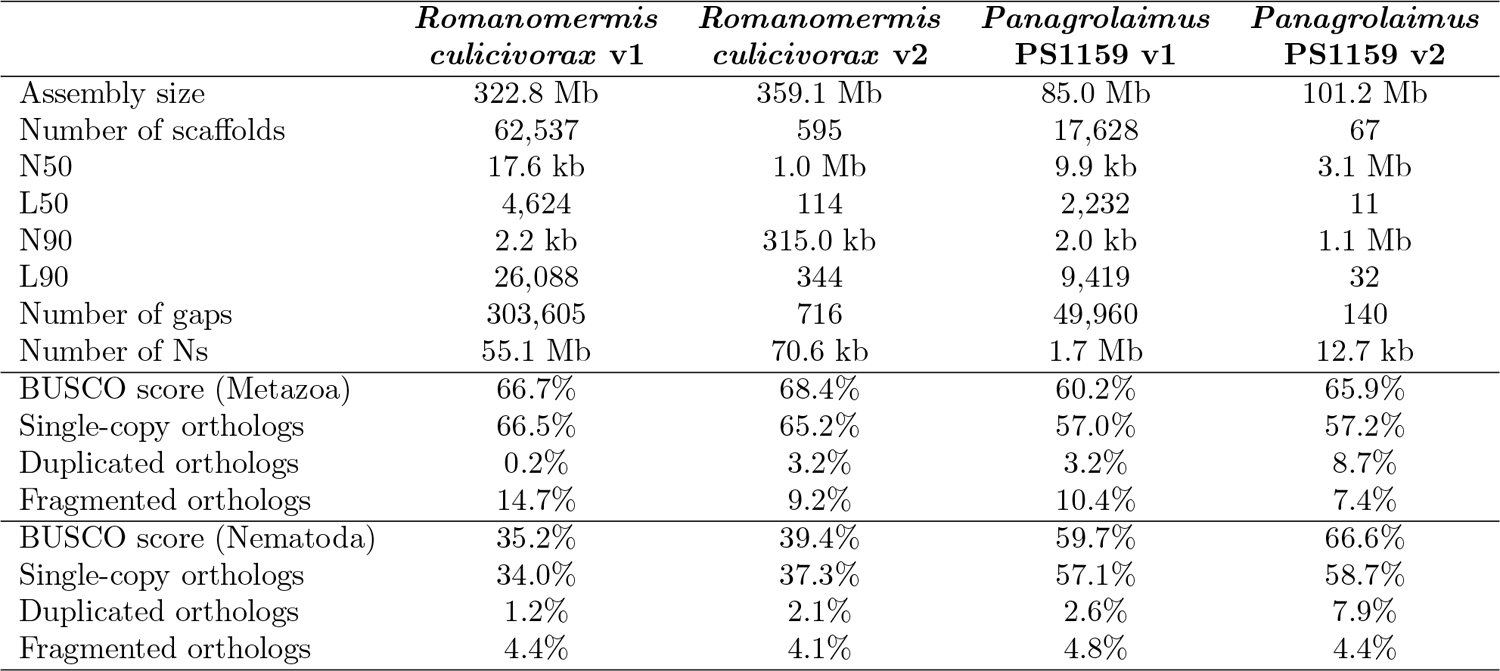
Assembly statistics of previous (v1) and new (v2) versions of *Romanomermis culicivorax* and *Panagrolaimus* sp. PS1159.

### 3.3 Repeat and gene annotation

Repetitions were better resolved in the new long-read assemblies than in the originally published ones (Figure 3). 68.2% of repeats were identified in the assembly of *R. culicivorax*, bringing it closer to the repetitive content of *Mermis negrescens*. The assembly of *P.* sp. PS1159 only has 16.0%, which is also higher than the 7.2% of repeats in the Illumina assembly. Many transposable elements (TE) were recovered in these improved assemblies that were undetected in Illumina assemblies. Notably, more long terminal repeats (LTR) were identified in *R. culicivorax*, the number of target inverted repeats was greatly increased, and 6.5 Mb of polintons were uncovered while they were almost absent in the Illumina assembly (Table S3). The load of transposable elements is much lower in *P.* sp. PS1159 but still has a wider variety of LTRs, TIRs, helitrons and other elements than the Illumina assembly. Gene prediction resulted in 16,689 annotated genes for *R. culicivorax*, with overall BUSCO scores of 77.6% (Metazoa) and 56.2% (Nematoda), and 27,203 annotated genes for *P.* sp. PS1159 with overall BUSCO scores of 77.3% (Metazoa) and 78.6% (Nematoda). As expected, these annotations are more complete than the ones published with the previous Illumina assemblies.

**Figure 3:**
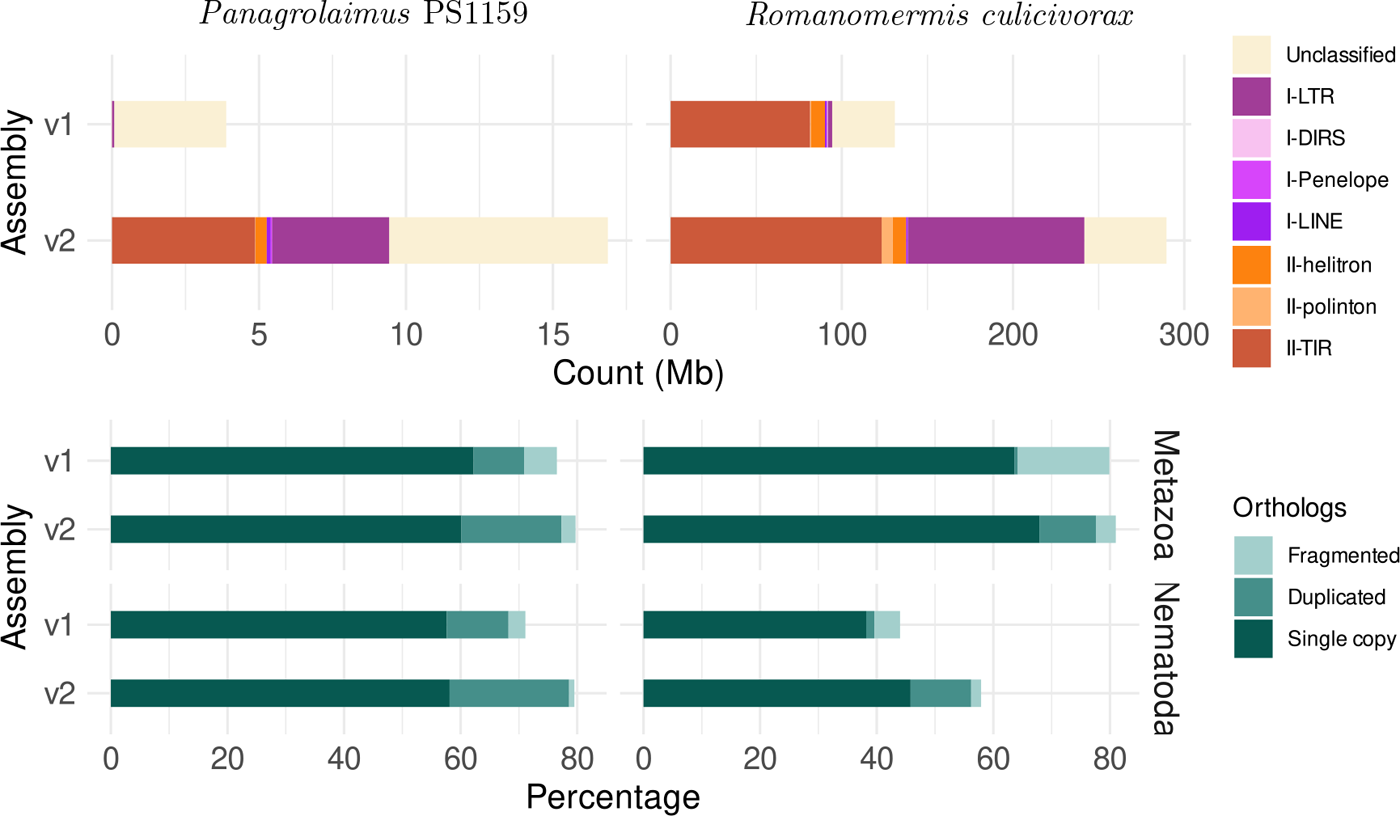
Comparison of assemblies based on TE (top) count and BUSCO ortholog (bottom) statistics for the protein annotations (against the Metazoa and Nematoda lineages) shows higher repeat and gene completeness of the new assemblies.

### 3.4 Orthology analyses of the phased assembly of *Panagrolaimus* sp. PS1159

To re-analyse the *Panagrolaimus* sp. PS1159 in regard to being a triploid genome, we selected the hybrid hifiasm assembly as a phased candidate. After decontamination, this assembly has a size of 264.7 Mb, 876 contigs and an N50 of 559 kb. The assembly’s BUSCO scores have high numbers of duplicated orthologs: 1.8% single-copy orthologs and 65.3% duplicated orthologs against Metazoa; 2.1% single-copy orthologs and 66.3% duplicated orthologs against Nematoda. The *k* -mer spectrum shows that the assembly has *k* -mers represented once, twice, or in three copies in the three different peaks at 50X, 100X and 150X (Figure S7), which is expected for a phased triploid genome assembly. In addition, the QV score reaches 48.18. Annotation resulted in 70,448 predicted genes, with BUSCO scores of 78.1% (77.4% duplicated) against Metazoa and 79.7% (78.7% duplicated) against Nematoda. We analyzed the number of ortholog copies from the annotated genes in the collapsed and phased assemblies (Figure 4), considering that orthologs used by BUSCO are expected as single copy. For the collapsed assembly, most orthologs are in only one copy. In the phased assembly, the majority of orthologs are in three copies, as there would be one copy for each haplotype. This brings further support to the triploidy of *Panagrolaimus* sp. PS1159.

**Figure 4:**
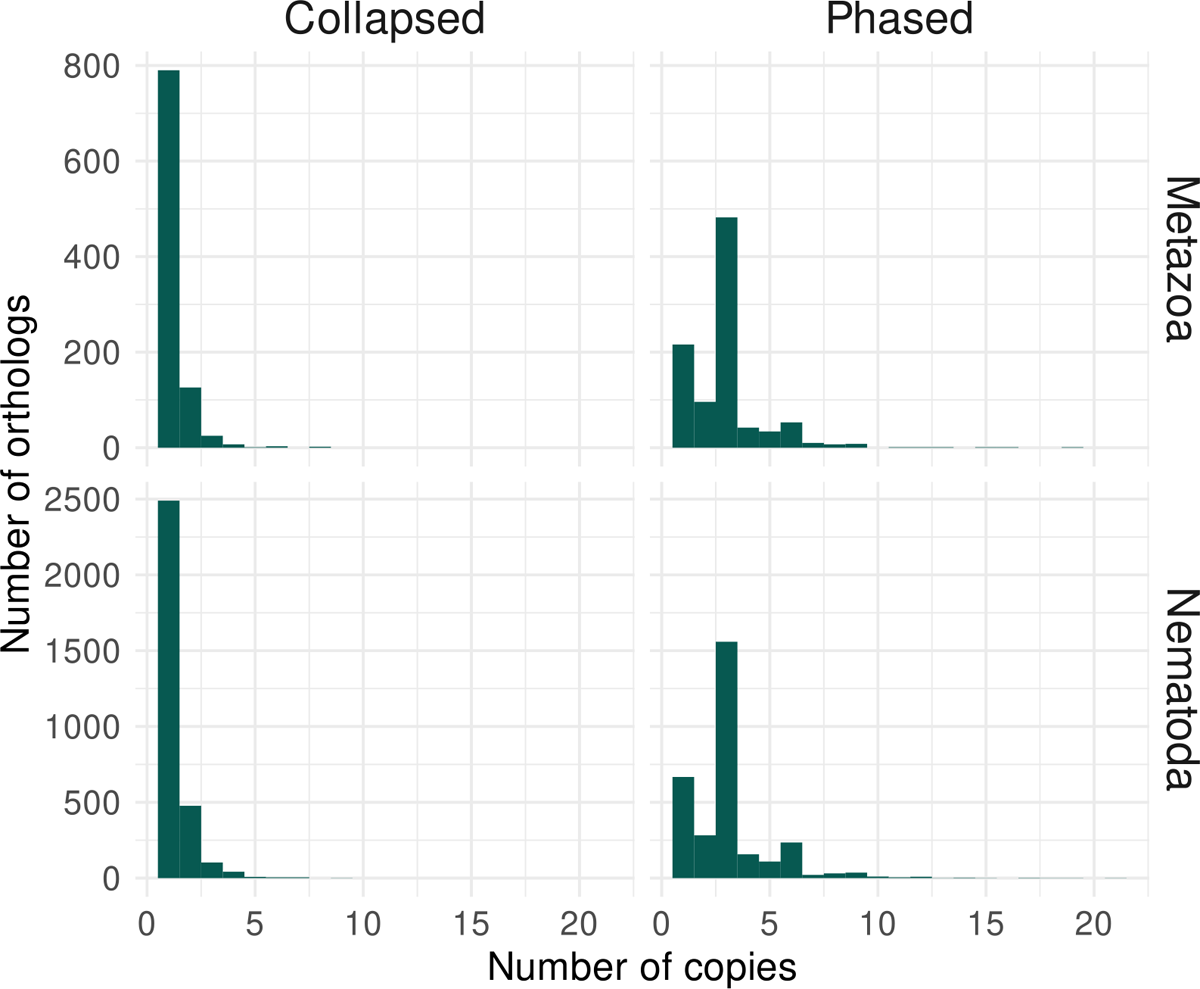
Ortholog analysis of *Panagrolaimus* sp. PS1159 collapsed and phased assemblies supports triploidy. The histograms represent the number of orthologs with their copy number from the lineages Metazoa and Nematoda identified in the protein-coding genes annotated for *P.* sp. PS1159. In the collapsed assembly, the majority of orthologs are present in a single copy, while most orthologs are in three copies in the phased assembly.

## 4 Discussion

Our new long-read assemblies for *Romanomermis culicivorax* and *Panagrolaimus* sp. PS1159 provide a drastic improvement to the previously published short-read-based assemblies, with higher contiguity, improved repeat resolution, and more accurate gene annotation. Furthermore, we generated a draft phased assembly of *P.* sp. PS1159, which opens new possibilites for haplotype-specicific analyses. With the addition of long-range sequencing data, such as chromosome conformation capture, we can expect to scaffold these high-quality assemblies into chromosome candidates and further investigate genome structures.

The first challenge consisted in generating PacBio HiFi and Nanopore sequencing data for these two nonmodel species. The resulting reads clearly highlight the strengths of these technologies: while Nanopore reads provide an advantage on length, PacBio HiFi reads have the highest accuracy. It should be noted however that the overall accuracy of Nanopore reads has increased compared to data from R9.4.1 flowcells [52] and were sufficient to produce assemblies with a high BUSCO completeness. Ultra-low input PacBio HiFi sequencing resulted in large datasets (over 29 Gb) despite the use of only a few individuals, and also led to high-quality draft assemblies. This amplification-based approach can be favored when the DNA availability for a species is limited, and for instance for nematodes which cannot be cultured.

Most initial assemblies improved on the published Illumina assemblies of the two species. The oversized PacBio HiFi assemblies of *R. culicivorax* could be attributed to the use of several individuals combined with the high accuracy of PacBio HiFi reads, leading to the separation of multiple haplotypes in heterozygous regions. As Nanopore reads had a lower accuracy, alternative haplotypes would not have been discriminated from errors and did not result in such large assemblies. For *P.* sp. PS1159, the Nanopore dataset was large enough to select for the more accurate Q20+ reads; therefore, haplotypes could be separated in both PacBio HiFi and Nanopore assemblies. In fact, almost all draft assemblies had the three haplotypes mostly separated with sizes close to 300 Mb (which would be the expected phased assembly size) and most BUSCO orthologs in multiple copies. Regarding contiguity, *R. culicivorax* assemblies were generally less contiguous than *P.* sp. PS1159 assemblies, which might be attributed to the higher repetitive content of this genome.

The most striking improvement in these assemblies lies in the resolution of repetitive regions. For both species, the percentage of repetitions in the genomes increased and revealed a wider variety of transposable elements. The comparison highlights that these transposable elements were in fact almost absent in the assembly of *P.* sp. PS1159 and very partially recovered in the assembly of *R. culicivorax*. Considering that TEs represent 289 Mb of the 359-Mb genome, we can estimate that a large aspect of this genome was completely overlooked in the past. A recent study has shown that genome assemblies from basal nematodes contain more repeats (ranging from 23.4% up to 50.6% repeats) than nematodes belonging to other clades (ranging from 0.8 %p to 31%) [53]. The results here presented are consistent with the previousfindings as *R. culicivorax*, a basal nematode, showed a high repeat and TE content and the derived *P.* sp. PS1159 has a low repeat and TE content. These variations and the better resolution of repetitions in long-read assembly should prompt further investigation into TE contents through nematode evolution.

The use of high-accuracy long reads permitted the generation of a first draft phased assembly of *P.* sp. PS1159. This assembly, combined with *k* -mer predictions based on PacBio HiFi reads and the analysis of Schiffer et al. [19], confirms that this species has a triploid genome. Considering the potential hybridization which could have introduced this third copy, a haplotype-resolved assembly is especially warranted to identify the original and newly acquired alleles. These analyses demonstrate the feasibility of long-read collapsed and phased assemblies for challenging genomes of understudied nematode species, including in the context of high repetitiveness and polyploidy. We gained new insights into these genomes regarding their gene and repeat content, which pave the way for more in-depth comparative genomics.

## Supplementary information

Supplementary figures and tables are available in Supplementary material. Command lines are available at github.com/worm-lab/Revisiting genomes.

## Data availability

The data for this study have been deposited in the European Nucleotide Archive (ENA) at EMBL-EBI under accession number PRJEB66727.

## Acknowledgements

We thank Christopher Kraus for his contribution to RNA sequencing and the Cologne Center for Genomics and the Genomics and Transcriptomics Laboratory for generation of sequencing data.

## References

[1] Rice, E. S. & Green, R. E. New approaches for genome assembly and scaffolding. Annual Review of Animal Biosciences7, 17–40 (2019).

[2] Wenger, A. M. et al. Accurate circular consensus long-read sequencing improves variant detection and assembly of a human genome. Nature Biotechnology37, 1155—-1162 (2019).

[3] Sereika, M. et al. Oxford Nanopore R10.4 long-read sequencing enables near-perfect bacterial genomes from pure cultures and metagenomes without short-read or reference polishing. bioRxiv(2021).

[4] Guiglielmoni, N., Rivera-Vicéns, R., Koszul, R. & Flot, J.-F. A deep dive into genome assemblies of non-vertebrate animals. Peer Community Journal2(2022).

[5] Shahid, S. & Slotkin, R. K. The current revolution in transposable element biology enabled by long reads. Current Opinion in Plant Biology54, 49–56 (2020).

[6] Cheng, H., Concepcion, G. T., Feng, X., Zhang, H. & Li, H. Haplotype-resolvedde novoassembly using phased assembly graphs with hifiasm. Nature Methods18, 1–6 (2021).

[7] Rautiainen, M. et al. Telomere-to-telomere assembly of diploid chromosomes with Verkko. Nature Biotechnology1–9 (2023).

[8] Hodda, M. Phylum nematoda: trends in species descriptions, the documentation of diversity, systematics, and the species concept. Zootaxa5114, 290–317 (2022).

[9] Kumar, S., Schiffer, P. H. & Blaxter, M. 959 nematode genomes: a semantic wiki for coordinating sequencing projects. Nucleic Acids Research40, D1295–D1300 (2012).

[10] Kumar, S., Koutsovoulos, G., Kaur, G. & Blaxter, M. Toward 959 nematode genomes. In Worm, vol. 1, 42–50 (2012).

[11] Presswell, B., Evans, S., Poulin, R. & Jorge, F. Morphological and molecular characterization of Mer-misnigrescens Dujardin, 1842 (Nematoda: Mermithidae) parasitizing the introduced European earwig (Dermaptera: Forficulidae) in New Zealand. Journal of Helminthology89, 267–276 (2015).

[12] Giblin, R. M. & Platzer, E. G. Romanomermis culicivorax parasitism and the development, growth, and feeding rates of two mosquito species. Journal of Invertebrate Pathology46, 11–19 (1985).

[13] Petersen, J., Chapman, H., Willis, O. & Fukuda, T. Release ofRomanomermis culicivoraxfor the control ofAnopheles albimanusin El Salvador II. Application of the nematode. The American Journal of Tropical Medicine and Hygiene27, 1268–1273 (1978).

[14] Abagli, A. Z., Alavo, T. B., Perez-Pacheco, R. & Platzer, E. G. Efficacy of the mermithid nematode, Romanomermis iyengari, for the biocontrol of Anopheles gambiae, the major malaria vector in sub-saharan africa. Parasites & Vectors12, 1–8 (2019).

[15] Mitreva, M. et al. The draft genome of the parasitic nematode Trichinella spiralis. Nature Genetics43, 228–235 (2011).

[16] Bhattarai, U. R., Poulin, R., Gemmell, N. J. & Dowle, E. Genome assembly and annotation of the mermithid nematode Mermis nigrescens. bioRxiv2022–11 (2022).

[17] Schiffer, P. H. et al. The genome of Romanomermis culicivorax: revealing fundamental changes in the core developmental genetic toolkit in Nematoda. BMC Genomics14, 1–16 (2013).

[18] Lewis, S. C. et al. Molecular evolution inPanagrolaimusnematodes: origins of parthenogenesis, hermaphroditism and the Antarctic speciesP. davidi. BMC Evolutionary Biology9(2009).

[19] Schiffer, P. H. et al. Signatures of the Evolution of Parthenogenesis and Cryptobiosis in the Genomes of Panagrolaimid Nematodes. iScience21, 587–602 (2019).

[20] Shatilovich, A. et al. A novel nematode species from the siberian permafrost shares adaptive mechanisms for cryptobiotic survival withC. elegansdauer larva. PLOS Genetics19, e1010798 (2023).

[21] McGill, L. M. et al. Anhydrobiosis and freezing-tolerance: Adaptations that facilitate the establishment of panagrolaimus nematodes in polar habitats. PLOS ONE10, e0116084 (2015).

[22] Shannon, A. J., Browne, J. A., Boyd, J., Fitzpatrick, D. A. & Burnell, A. M. The anhydrobiotic potential and molecular phylogenetics of species and strains of Panagrolaimus(Nematoda, Panagrolaimidae). Journal of Experimental Biology208, 2433–2445 (2005).

[23] Koetsier, P. A. G. & Cantor, E. J. A simple approach for effective shearing and reliable concentration measurement of ultra-high-molecular-weight DNA. BioTechniques71, 439–444 (2021).

[24] Oxford Nanopore Technologies. Dorado,https://github.com/nanoporetech/dorado (2022).

[25] Danecek, P. et al. Twelve years of SAMtools and BCFtools. GigaScience10(2021). Giab008.

[26] De Coster, W. & Rademakers, R. NanoPack2: population-scale evaluation of long-read sequencing data. Bioinformatics39, btad311 (2023).

[27] Chomczynski, P. & Sacchi, N. Single-step method of RNA isolation by acid guanidinium thiocyanate-phenol-chloroform extraction. Analytical Biochemistry162, 156–159 (1987).

[28] Ranallo-Benavidez, T. R., Jaron, K. S. & Schatz, M. C. GenomeScope 2.0 and Smudgeplot for reference-free profiling of polyploid genomes. Nature Communications11, 1432 (2020).

[29] Kolmogorov, M., Yuan, J., Lin, Y. & Pevzner, P. A. Assembly of long, error-prone reads using repeat graphs. Nature Biotechnology37, 540–546 (2019).

[30] NextOmics. NextDenovo,https://github.com/Nextomics/NextDenovo (2019).

[31] Ruan, J. & Li, H. Fast and accurate long-read assembly with wtdbg2. Nature Methods17, 155–158 (2020).

[32] Koren, S. et al. Canu: scalable and accurate long-read assembly via adaptivek-mer weighting and repeat separation. Genome Research25, 1–11 (2017).071282.

[33] Li, H. seqtk, https://github.com/lh3/seqtk (2012).

[34] Pathogen Informatics, Wellcome Sanger Institute. assembly-stats,https://github.com/sanger-pathogens (2014).

[35] Manni, Mosè and Berkeley, Matthew R and Seppey, Mathieu and Simão, Felipe A and Zdobnov, Evgeny M. BUSCO update: novel and streamlined workflows along with broader and deeper phylogenetic coverage for scoring of eukaryotic, prokaryotic, and viral genomes. Molecular Biology and Evolution38, 4647–4654 (2021).

[36] Li, H. Minimap2: pairwise alignment for nucleotide sequences. Bioinformatics34, 3094–3100 (2018).

[37] Altschul, S. F., Gish, W., Miller, W., Myers, E. W. & Lipman, D. J. Basic local alignment search tool. Journal of Molecular Biology215, 403–410 (1990).

[38] Challis, R., Richards, E., Rajan, J., Cochrane, G. & Blaxter, M. Blobtoolkit–interactive quality assessment of genome assemblies. G3: Genes, Genomes, Genetics10, 1361–1374 (2020).

[39] Guan, D. et al. Identifying and removing haplotypic duplication in primary genome assemblies. Bioinformatics36, 2896–2898 (2020).

[40] Alonge, M. et al. Automated assembly scaffolding using RagTag elevates a new tomato system for high-throughput genome editing. Genome Biology23, 1–19 (2022).

[41] Ou, S. et al. Benchmarking transposable element annotation methods for creation of a streamlined, comprehensive pipeline. Genome Biology20, 1–18 (2019).

[42] Gremme, G., Steinbiss, S. & Kurtz, S. GenomeTools: a comprehensive software library for efficient processing of structured genome annotations. IEEE/ACM Transactions on Computational Biology and Bioinformatics 10, 645–656 (2013).

[43] Xu, Z. & Wang, H. LTR FINDER: an efficient tool for the prediction of full-length LTR retrotransposons. Nucleic Acids Research35, W265–W268 (2007).

[44] Ou, S. & Jiang, N. LTR retriever: a highly accurate and sensitive program for identification of long terminal repeat retrotransposons. Plant Physiology176, 1410–1422 (2018).

[45] Xiong, W., He, L., Lai, J., Dooner, H. K. & Du, C. HelitronScanner uncovers a large overlooked cache of Helitron transposons in many plant genomes. Proceedings of the National Academy of Sciences111, 10263–10268 (2014).

[46] Shi, J. & Liang, C. Generic Repeat Finder: a high-sensitivity tool for genome-widede novorepeat detection. Plant Physiology180, 1803–1815 (2019).

[47] Su, W., Gu, X. & Peterson, T. TIR-Learner, a new ensemble method for TIR transposable element annotation, provides evidence for abundant new transposable elements in the maize genome. Molecular Plant12, 447–460 (2019).

[48] Flynn, J. M. et al. RepeatModeler2 for automated genomic discovery of transposable element families. Proceedings of the National Academy of Sciences117, 9451–9457 (2020).

[49] Kim, D., Paggi, J. M., Park, C., Bennett, C. & Salzberg, S. L. Graph-based genome alignment and genotyping with HISAT2 and HISAT-genotype. Nature Biotechnology37, 907–915 (2019).

[50] Gabriel, L. et al. BRAKER3: Fully Automated Genome Annotation Using RNA-Seq and Protein Evidence with GeneMark-ETP, AUGUSTUS and TSEBRA. bioRxiv2023–06 (2023).

[51] Rhie, A., Walenz, B. P., Koren, S. & Phillippy, A. M. Merqury: reference-free quality, completeness, and phasing assessment for genome assemblies. Genome Biology21, 1–27 (2020).

[52] Guiglielmoni, N., Houtain, A., Derzelle, A., Van Doninck, K. & Flot, J.-F. Overcoming uncollapsed haplotypes in long-read assemblies of non-model organisms. BMC Bioinformatics22, 1–23 (2021).

[53] Lee, Y.-C. et al. Single-worm long-read sequencing reveals genome diversity in free-living nematodes. Nucleic Acids Research51, 8035–8047 (2023).

