## Supplementary_material for "Revisiting genomes of non-model species with long reads yields new insights into their biology and evolution"

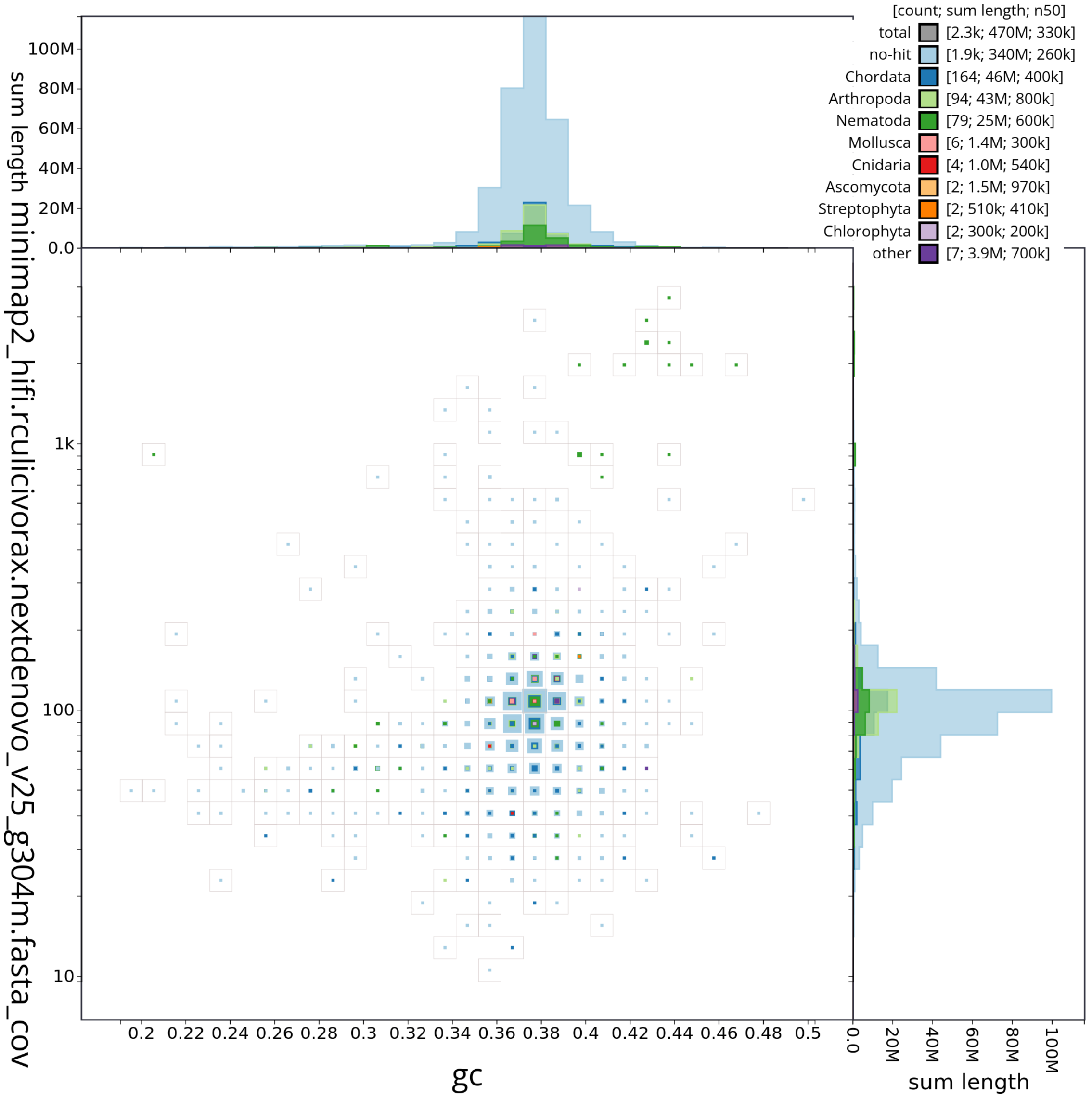


Figure S1: Example of Blobtools analysis for the NextDenovo PacBio HiFi assembly of *Romanomermis culicivorax*.

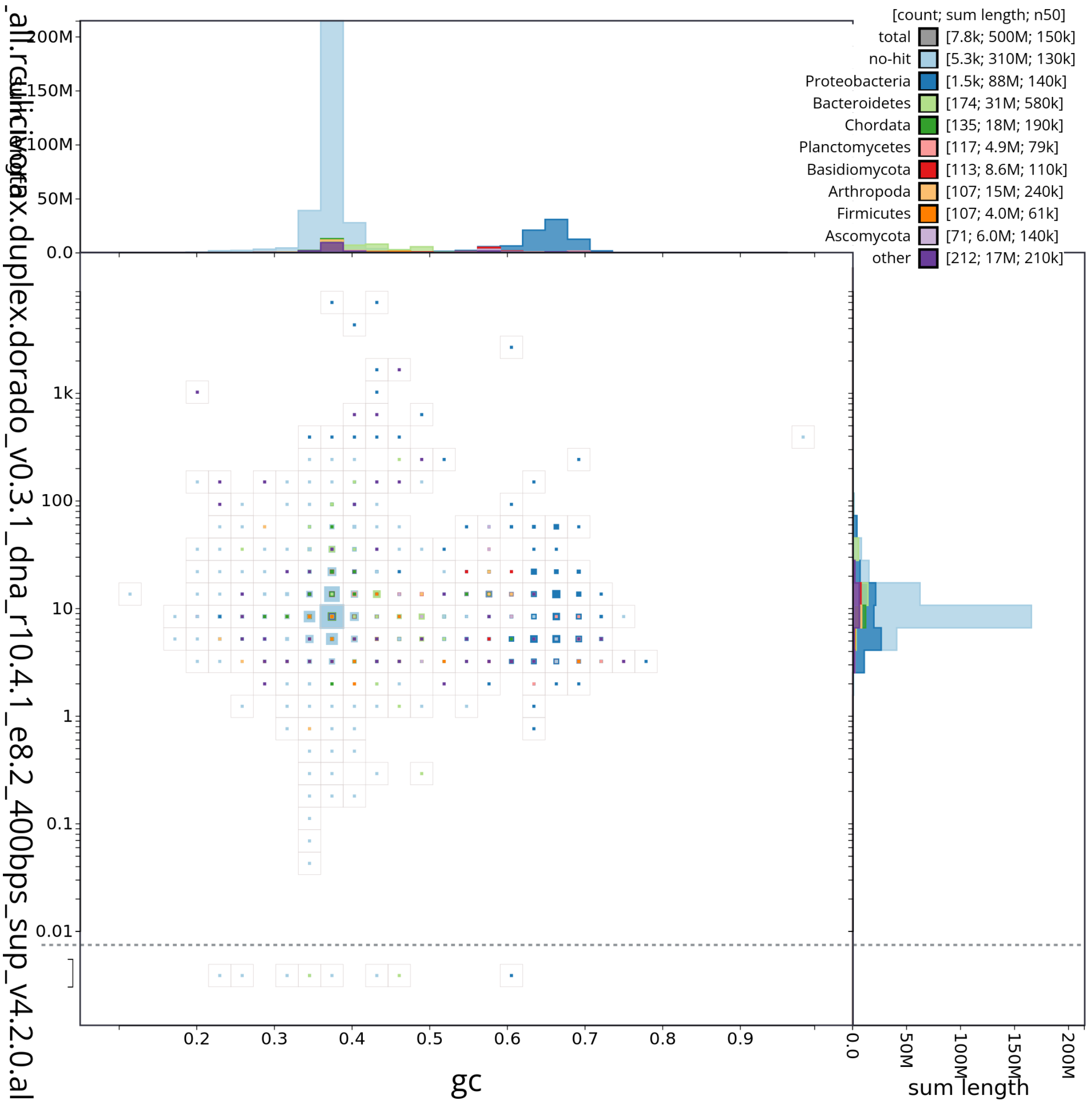


Figure S2: Example of Blobtools analysis for the Flye Nanopore assembly of *Romanomermis culicivorax*.

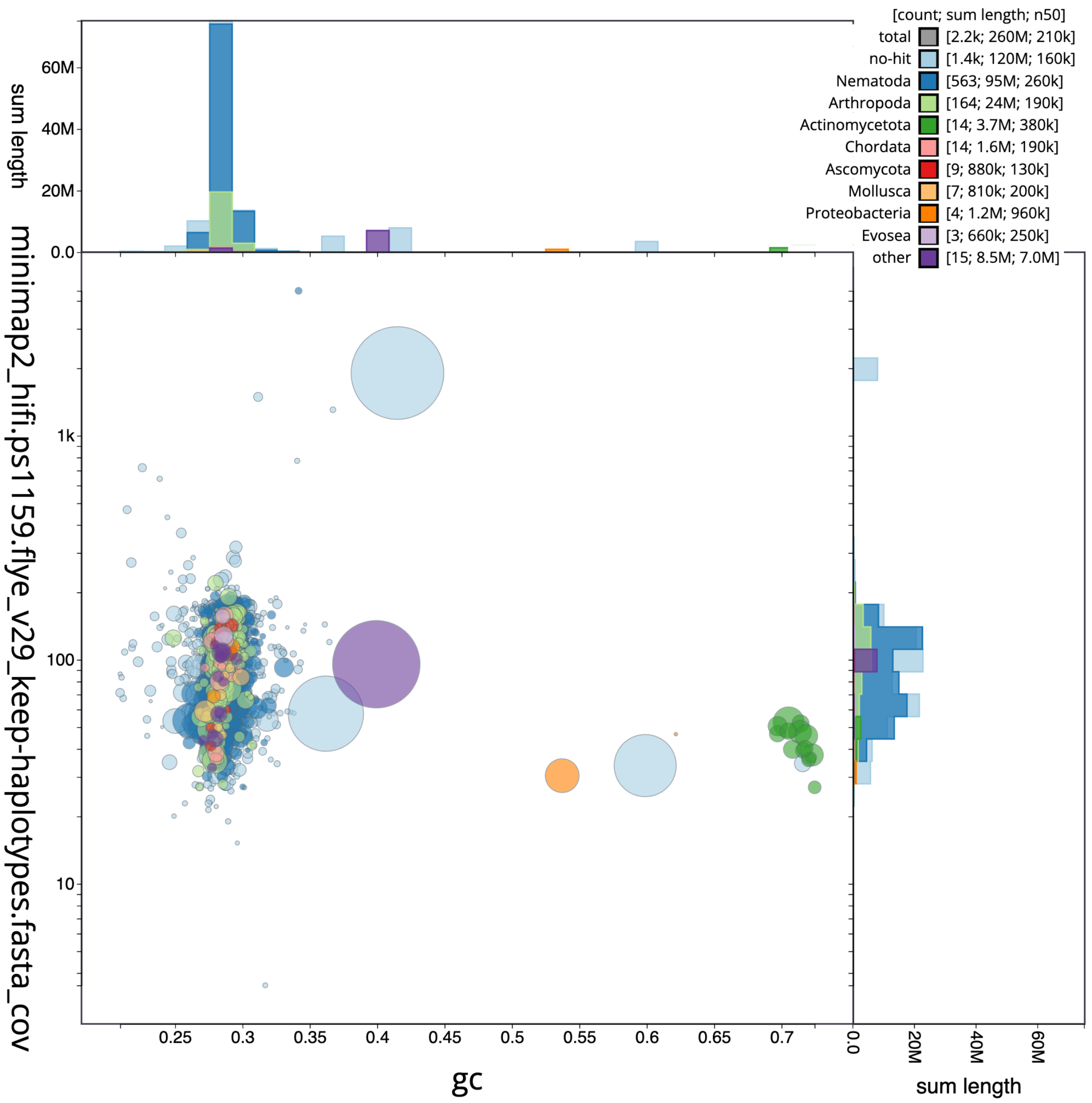


Figure S3: Example of Blobtools analysis for the Flye Nanopore assembly of *Panagrolaimus* sp. PS1159.

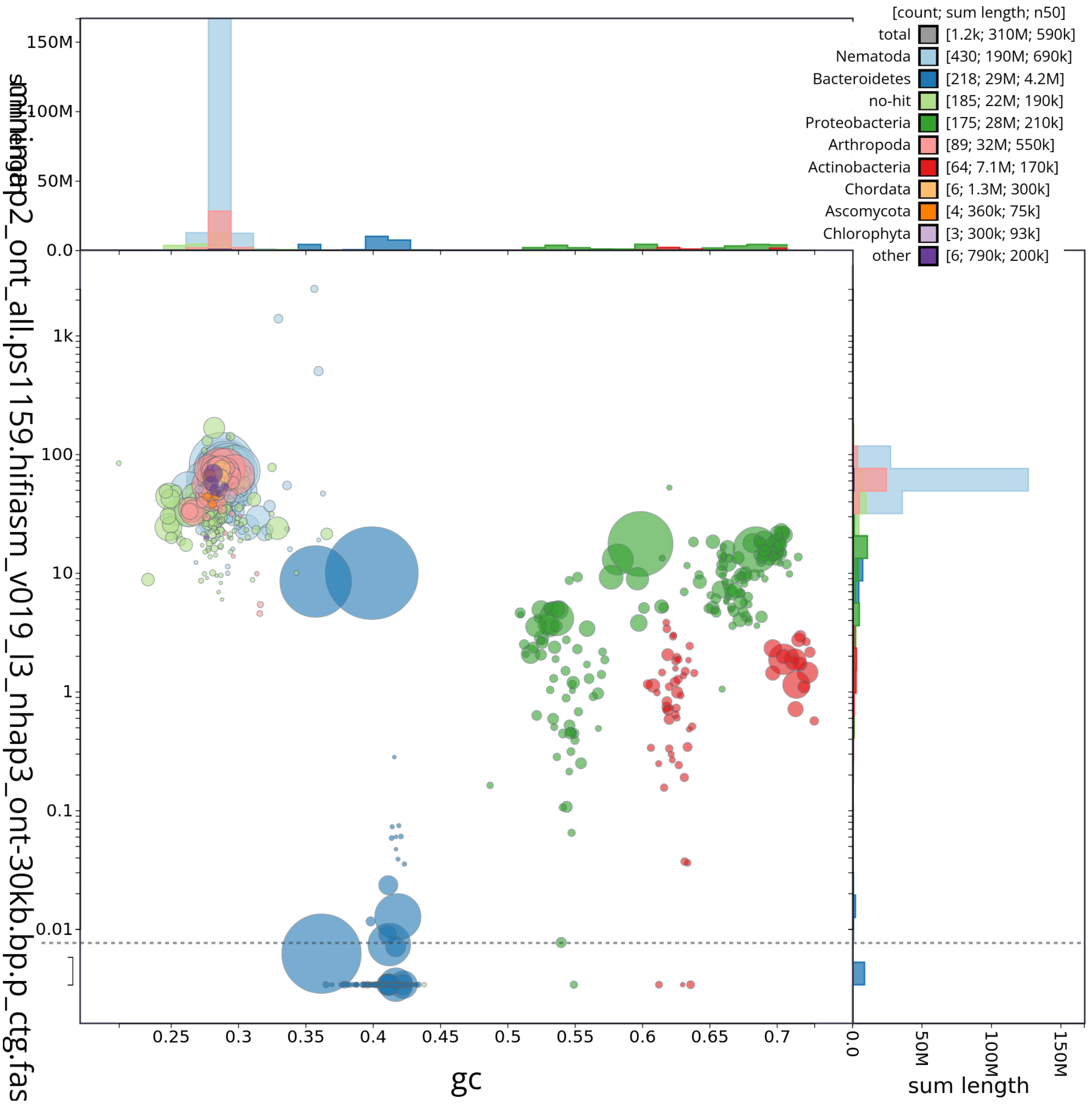


Figure S4: Example of Blobtools analysis for the hifiasm PacBio HiFi + Nanopore assembly of *Panagrolaimus* sp. PS1159.

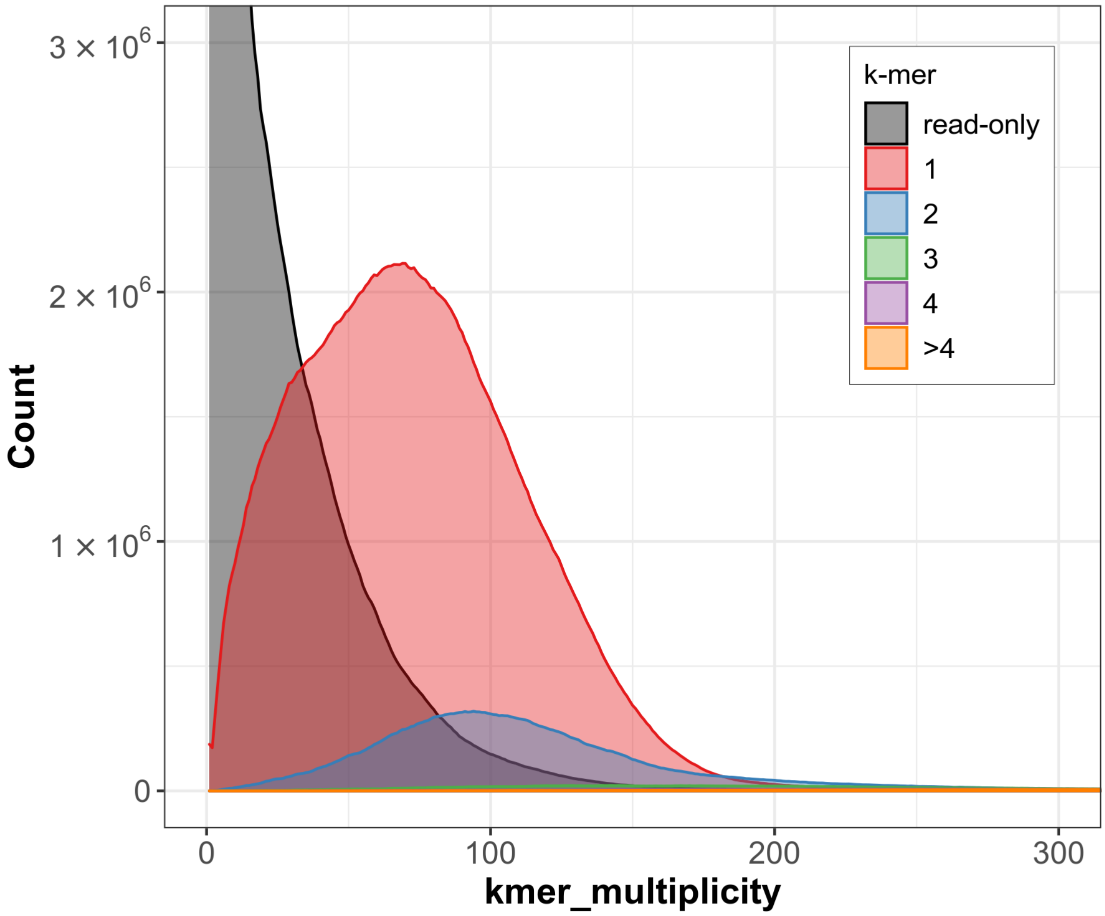


Figure S5: Merqury analysis of the final assembly of *Romanomermis culicivorax*.

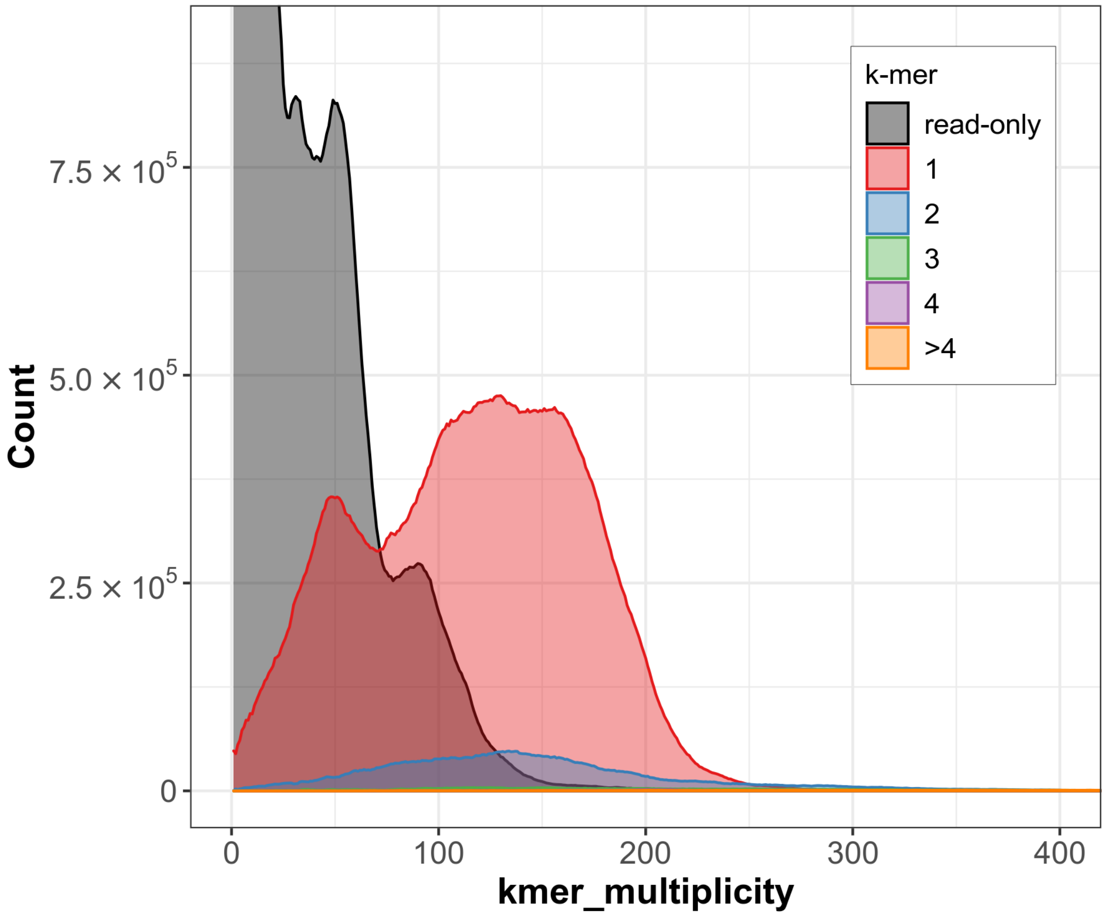


Figure S6: Merqury analysis of the final assembly of *Panagrolaimus* sp. PS1159.

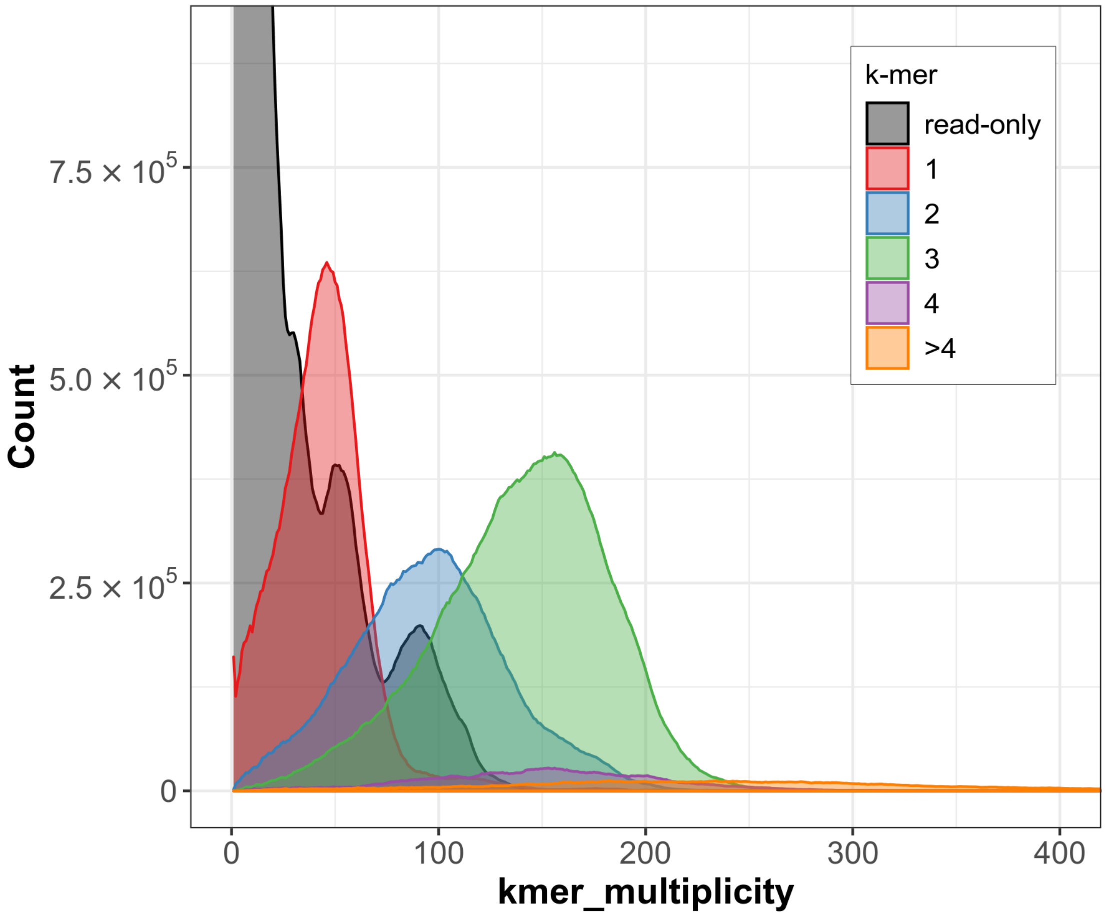


Figure S7: Merqury analysis of the phased assembly of *Panagrolaimus* sp. PS1159.

Table S1: Statistics of the PacBio HiFi datasets.

| **Species** | **Size** | **N50** |
|---|---|---|
| *Romanomermis culicivorax* | 37,542,673,078 | 12,613 |
| *Panagrolaimus* sp. PS1159 | 29,159,296,422 | 15,797 |

Table S2: Statistics of the Nanopore datasets.

| **Species** | **Quality threshold** | **Size** | **N50** | **Largest** |
|---|---|---|---|---|
| *Romanomermis culicivorax* | 0 | 5,697,172,871 | 15,936 | 540,992 |
| | 10 | 5,021,948,640 | 15,980 | 218,858 |
| | 15 | 3,943,100,749 | 15,551 | 218,858 |
| | 20 | 979,904,894 | 15,886 | 115,632 |
| *Panagrolaimus* sp. PS1159 | 0 | 10,696,877,601 | 33,428 | 914,599 |
| | 10 | 9,344,589,639 | 33,814 | 205,535 |
| | 15 | 8,280,860,091 | 33,820 | 205,535 |
| | 20 | 4,718,164,203 | 34,667 | 173,662 |

| Species | Assembly | Category | Count |
|---|---|---|---|
| *Romanomermis culicivorax* | v1 | Unclassified | 36,611,241 |
| | | I-LTR | 2,623,729 |
| | | I-DIRS | 380,102 |
| | | I-Penelope | 198,523 |
| | | I-LINE | 987,323 |
| | | II-helitron | 8,326,702 |
| | | II-polinton | 444,560 |
| | | II-TIR | 81,385,493 |
| | v2 | Unclassified | 47,999,208 |
| | | I-LTR | 103,047,244 |
| | | I-DIRS | 71,775 |
| | | I-Penelope | 0 |
| | | I-LINE | 881,202 |
| | | II-helitron | 7,860,451 |
| | | II-polinton | 6,478,667 |
| | | II-TIR | 123,251,468 |
| *Panagrolaimus* sp. PS1159 | v1 | Unclassified | 3,813,266 |
| | | I-LTR | 59,899 |
| | | I-DIRS | 0 |
| | | I-Penelope | 0 |
| | | I-LINE | 6,942 |
| | | II-helitron | 5,224 |
| | | II-polinton | 0 |
| | | II-TIR | 475 |
| | v2 | Unclassified | 7,442,799 |
| | | I-LTR | 3,983,941 |
| | | I-DIRS | 0 |
| | | I-Penelope | 143,146 |
| | | I-LINE | 44,333 |
| | | II-helitron | 380,276 |
| | | II-polinton | 18,632 |
| | | II-TIR | 4,864,827 |

Table S3: Transposable elements counts.
